# *Mycobacterium marinum* phthiocerol dimycocerosates enhance macrophage phagosomal permeabilization and membrane damage

**DOI:** 10.1101/2020.05.04.076596

**Authors:** Morwan M. Osman, Antonio J. Pagán, Jonathan K. Shanahan, Lalita Ramakrishnan

## Abstract

Phthiocerol dimycocerosates (PDIMs) are a class of mycobacterial lipids that promote virulence in *Mycobacterium tuberculosis* and *Mycobacterium marinum*. It has recently been shown that PDIMs work in concert with the *M. tuberculosis* Type VII secretion system ESX-1 to permeabilize the phagosomal membranes of infected macrophages. As the zebrafish-*M. marinum* model of infection has revealed the critical role of PDIM at the host-pathogen interface, we set to determine if PDIMs contributed to phagosomal permeabilization in *M. marinum*. Using an *ΔmmpL7* mutant defective in PDIM transport, we find the PDIM-ESX-1 interaction to be conserved in an *M. marinum* macrophage infection model. However, we find PDIM and ESX-1 mutants differ in their degree of defect, with the PDIM mutant retaining more membrane damaging activity. Using an *in vitro* hemolysis assay— a common surrogate for cytolytic activity, we find that PDIM and ESX-1 differ in their contributions: the ESX-1 mutant loses hemolytic activity while PDIM retains it. Our observations confirm the involvement of PDIMs in phagosomal permeabilization in *M. marinum* infection and suggest that PDIM enhances the membrane disrupting activity of pathogenic mycobacteria and indicates that the role they play in damaging phagosomal and red blood cell membranes may differ.

## Introduction

*Mycobacterium tuberculosis*, the causative agent of tuberculosis (TB), is an obligate human pathogen; its ability to survive and spread is dependent upon its ability to survive within its human host. During the early stages of infection, this requires manipulation of the innate immune system, allowing for survival of the pathogen within the microbicidal environment of host macrophages[1]. *M. tuberculosis* and the pathogenic *M. marinum* use their lipid coats and Type VII secretion systems to evade and co-opt ordinarily lethal host defenses within these monocytes.

The virulence lipid phthiocerol dimycocerosate (PDIM) is an outer membrane virulence lipid in both *M. marinum* and *M. tuberculosis*. PDIM-deficient mutants in *M. marinum* and *M. tuberculosis* are attenuated in animal models of infection[2–4] and membrane permeability[5,6]. Contact between mycobacteria and host environment results in the transfer of surface PDIM into host membranes, suppressing toll-like receptor signaling (TLR) and preventing the recruitment of microbicidal monocytes during infection[4,7,8]. Consequently, PDIM mutants are rapidly phagocytosed and killed by microbicidal monocytes[4].

The ESX-1 Type VII secretion system was identified as essential for virulence when its loss was determined to be the primary cause of attenuation for the live BCG vaccine strain[9–12]. ESX-1 is crucial for survival within the macrophage during early infection, mediating evasion of bacterial killing and induction of growth-permissive responses. ESX-1 enhances recruitment of macrophages during infection, promoting granuloma formation[13–15]. Within the macrophage, ESX-1 mediates permeabilization of the mycobacterial phagosome[16,17]. Permeabilization induces the cGAS/STING and AIM2/NLRP3 cytosolic signaling pathways, which promote production of cytokines that may enhance bacterial survival[18–22].

Incorporation of PDIM into host membranes has been proposed to rigidify them, enhancing lysis by ESX-1[8]. Supporting this hypothesis, multiple groups have observed that loss of PDIM reduces phagosomal permeabilization in *M. tuberculosis*[23–26]. As the study of *M. marinum* has provided new insights into PDIM’s role in pathogenesis[4,7], we set out to determine if this PDIM-ESX-1 interaction was conserved in *M. marinum*. In this paper, we confirm that as in *M. tuberculosis*, proper PDIM localization is required for *M. marinum* to effectively permeabilize macrophage phagosomes.

## Results & discussion

ESX-1 is required for *M. marinum* and *M. tuberculosis* to permeabilize macrophage phagosomes during infection[16,17,27]. Recent work has shown that ESX-1 and PDIM are both required for permeabilization in *M. tuberculosis* [23–26]. We set out to determine if phagosomal permeabilization requires PDIM in *M. marinum* by using an attenuated *ΔmmpL7 M. marinum* mutant defective in PDIM localization to the mycomembrane[4]. To measure phagosomal permeabilization, we used the fluorescence resonance energy transfer (FRET)-based dye CCF4-AM as we have done previously[28]. Briefly, the lipophilic dye CCF4-AM is absorbed into the cytosol and is cleaved by cytosolic esterases. The resulting dye is retained in the cytosol and produces a green fluorescent signal (525 nm) upon excitation with a violet laser (405 nm)[17,29]. When phagosomal permeabilization occurs, the dye becomes accessible to *M. marinum*. *M. marinum* is capable of cleaving the dye with its endogenous mycobacterial β-lactamase BlaC, which causes a loss of FRET and an increase in blue fluorescence (450 nm). We used this dye to determine the relative ability of wildtype, *ΔESX-1*, and *ΔmmpL7 M. marinum* to permeabilize their phagosomes (Fig 1A-C). We found that both strains permeabilized their phagosomes less than wildtype *M. marinum* (Fig 1D). These results show that PDIM is required for *M. marinum* phagosomal permeabilization, as it is for *M. tuberculosis*.

**Figure 1:**
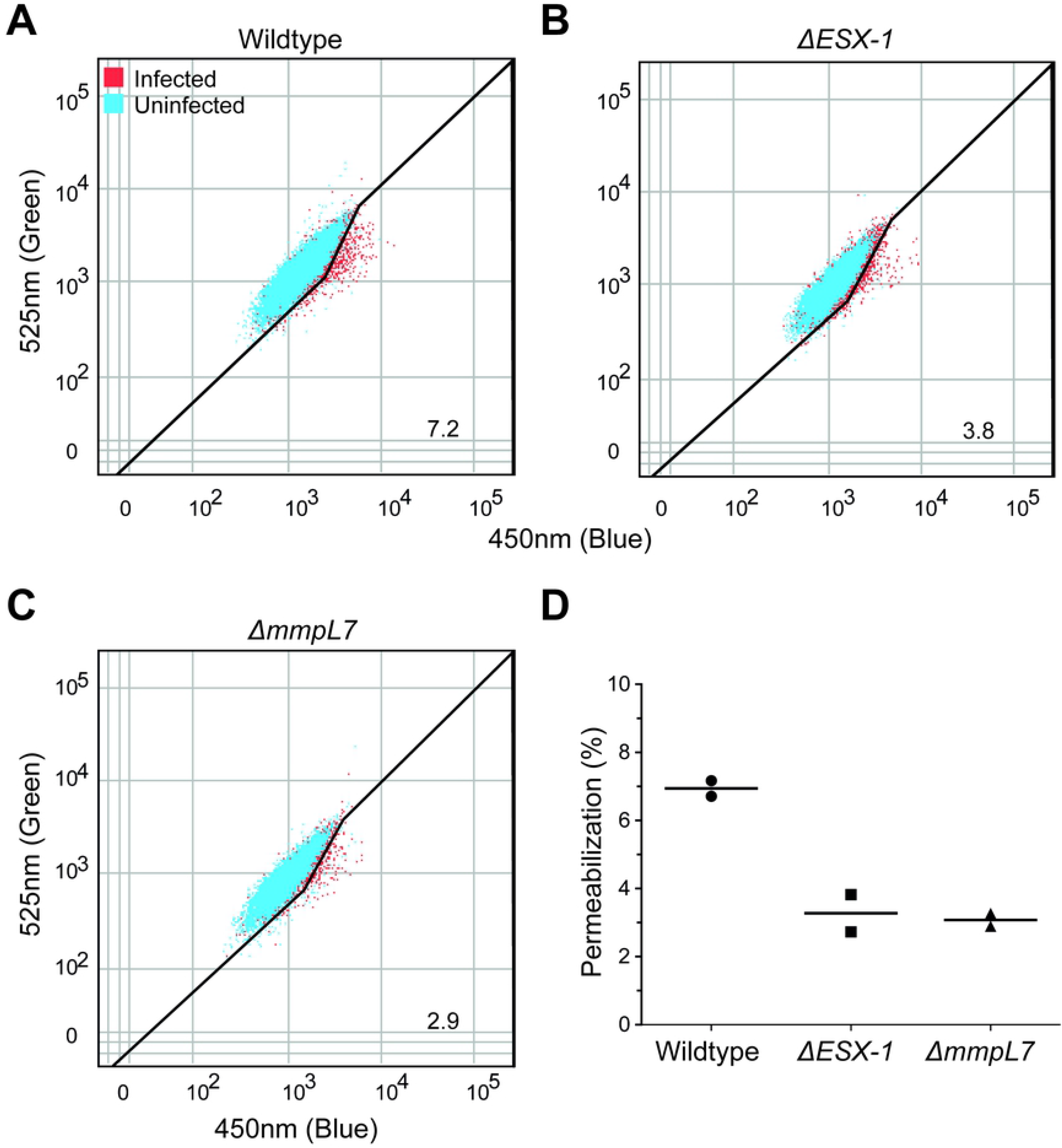
PDIM transport to the outer mycomembrane is required for optimal phagosomal permeabilization by *M. marinum*. Representative dot plots of (A) Wildtype, (B) Δ*ESX-1*, and (C) Δ*mmpL7* infected THP-1 macrophages 24 hours post infection with overlays of uninfected (blue) and infected (red) cells within each sample. Gating outlined in black, % permeabilization is as labeled. Plots are representative of two independent experiments. (D) Quantification of permeabilization events. Each point represents an independent experiment.

ESX-1-mediated phagosomal permeabilization is associated with membrane damage in both *M. tuberculosis* and *M. marinum* [16,17,27,28,30–32]. *M. tuberculosis* strains deficient in ESX-1 or PDIM synthesis show reduced galectin-3 or galectin-8 recruitment to intracellular sites of infection[23,24,33,34]. If the endosomal membrane is damaged, cytosolic galectins bind to lumenal β-galactoside-containing glycans that become exposed on damaged vesicles, and can be visualized by immunofluorescence microscopy[23,24,33–36]. Differing from the role of ESX-1 in phagosomal permeabilization during *M. tuberculosis* and *M. marinum* infection, the involvement of *M. marinum* PDIM in this process has not been demonstrated.

We next asked if *M. marinum* PDIM is similarly associated with phagosomal membrane damage. As expected, galectin-8 was readily observed at sites of infection with wildtype *M. marinum* and was greatly reduced with *ΔESX-1 M. marinum* (98% lower in *ΔESX-1*) (Fig 2). *ΔmmpL7 M. marinum* infection revealed an intriguing result: galectin-8 staining was reduced as compared to wildtype but less than *ΔESX-1* (84% lower in *ΔmmpL7*). Highlighting its intermediate level of staining, galectin-8 staining was seven times greater in *ΔmmpL7* than in *ΔESX-1* infection, (Fig 2). These results suggest that like in *M. tuberculosis*, *M. marinum* PDIM cooperates with ESX-1 to induce maximum damage to host membranes, and again PDIM surface localization is required.

**Figure 2.**
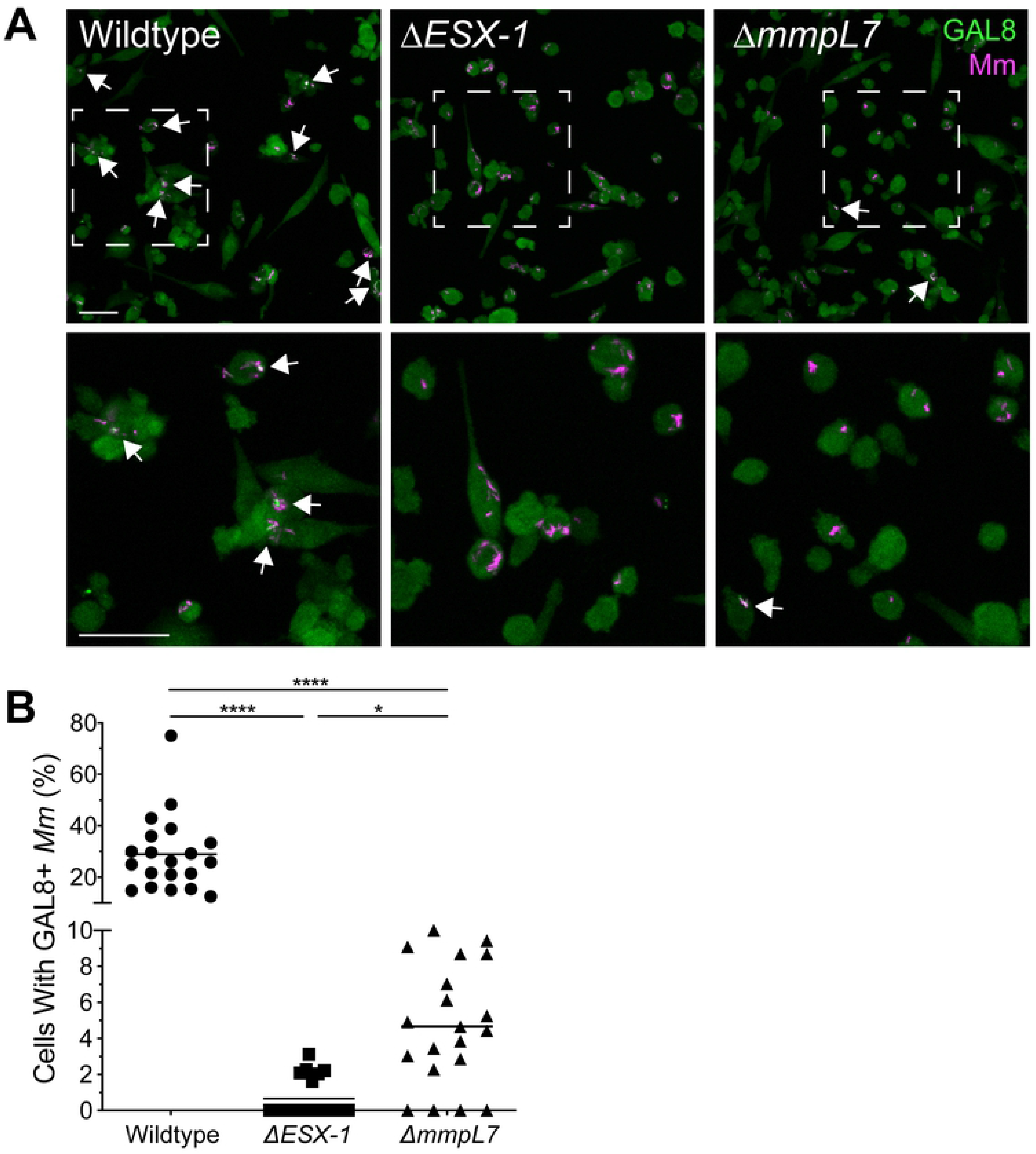
PDIM transport deficiency reduces *M. marinum*-induced phagosomal membrane damage. A. Maximum intensity projections of confocal micrographs showing galectin-8 labeling of PMA-differentiated THP-1 cells infected with tdTomato-expressing wildtype or mutant *M. marinum* strains 22 hours post infection. Arrows indicate co-localization of Galectin-8 (green) and *M. marinum* (magenta) fluorescence. Bottom panels show close-up views of the corresponding boxed regions in the top panels. Scale bars, 50μm. B. Percent of infected cells that have mycobacterial foci labeled with galectin-8. Symbols represent individual imaging fields. Horizontal lines depict mean values. The respective means and 95% confidence intervals for wildtype, *ΔESX-1* and *ΔmmpL7* infections are 28.9 (22.0 - 35.8), 0.7 (0.2 - 1.2), and 4.7 (3.1 - 6.3). Statistical significance was determined using a Kruskal-Wallis nonparametric test with Dunn’s post-hoc test to control for multiple comparisons, **** p < 0.0001, * p < 0.05. Data are representative of two experiments.

*M. tuberculosis* and *M. marinum* both show ESX-1 dependent hemolytic activity[37–39]. Furthermore, loss of hemolytic activity in an *M. marinum ΔESX-1* mutant can be rescued by complementing with the *M. tuberculosis* ESX-1 locus[28]. Historically, hemolysis in *M. marinum* has been regarded as a correlate of ESX-1 mediated virulence, although recent work suggests that hemolysis can be lost with minimal effects on intramacrophage growth or virulence[28,38,40–43]. As phagosomal permeabilization and hemolysis both require host membrane lysis, we set out to determine if the permeabilization-deficient *M. marinum ΔmmpL7* were defective in hemolysis as well. As expected, *ΔESX-1* had reduced hemolytic activity as compared to wildtype, *ΔmmpL7* had significantly more hemolytic activity than *ΔESX-1* (Fig 3). Thus, PDIM does not contribute to hemolysis as much as ESX-1, mirroring its lesser contribution to phagosomal membrane damage as evidenced by galectin-8 staining.

**Figure 3:**
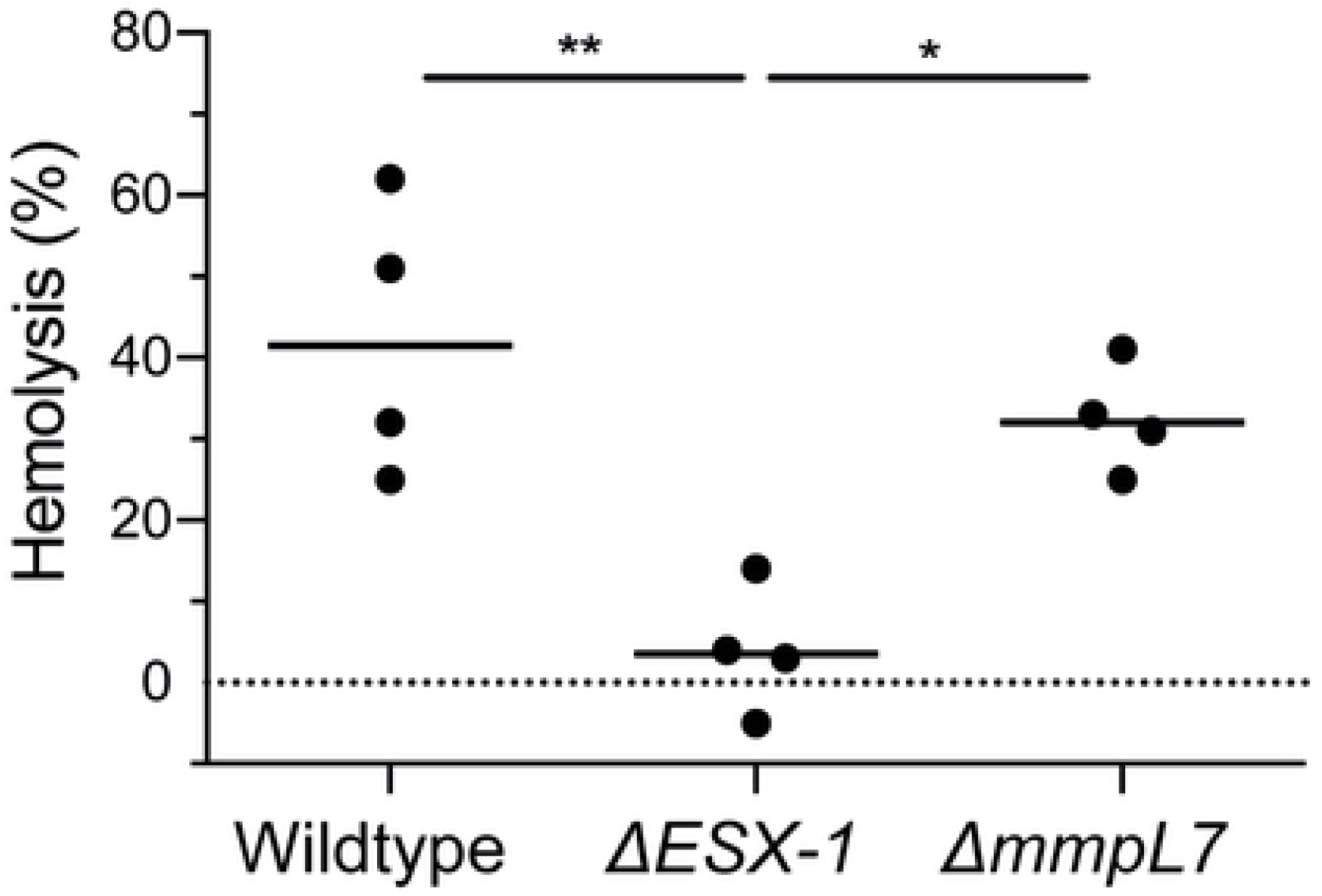
PDIM-deficient *M. marinum* retain hemolytic activity. 2-hour hemolysis of sheep red blood cells following incubation with Wildtype, Δ*ESX-1* and Δ*mmpL7 M. marinum* expressed as a percentage of complete detergent mediated lysis. Statistical significance was determined by one-way ANOVA with Tukey’s post-test, *p < 0.05, ** p < 0.01. Each point represents an independent experiment.

In summary, we find that proper PDIM localization enhances ESX-1 dependent phagosomal permeabilization in *M. marinum*. This matches observations that *M. tuberculosis* mutants in PDIM localization or synthesis are deficient in phagosomal permeabilization[23–26]. While we have not been able to directly determine the effect of PDIM loss on *M. marinum* ESX-1 secretion, the finding that *M. tuberculosis ΔmmpL7* retains its ESX-1 secretory activity [25] gives confidence that this is the case in *M. marinum* as well. Our finding that PDIM and ESX-1 contribute to the same extent to phagosomal membrane permeabilization when judged by the CCF4-AM assay, but differ in the galectin-8 phagosomal damage assay, suggests that the latter may be a more sensitive assay. Finally, the finding that *ΔmmpL7* is much more hemolytic than *ΔESX-1*, and nearly as hemolytic as wildtype, suggests that the hemolysis assay is not a reliable surrogate for phagosomal membrane damaging activity.

## Materials & methods

### Bacterial Strains

All strains were derived from wildtype *M. marinum* purchased from American Type Culture Collection (ATCC) (strain M, ATCC no. BAA-535). *M. marinum ΔESX-1* and *ΔmmpL7* were generated as described previously[4,14].

### Infections, galectin-8 immunofluorescence, and microscopy

2.5 × 10^5^ THP-1 cells were seeded on 24-well optical bottom tissue culture plates (Perkin Elmer, 1450-606), treated with 100nM phorbol 12-myrystate-13-acetate (PMA) (SIGMA, P1585) for two days. PMA-containing media was then removed and replaced with fresh media. Two days later, adherent cells were washed twice with PBS and infected with antibiotic-free media containing single-cell suspensions of tdTomato-expressing *M. marinum* at a multiplicity of infection of ~1.5 for 4 hours and maintained at 33°C, 5%CO_2_. 22 hours later, cells were fixed in 4% (wt/vol) paraformaldehyde in PBS at room temperature for at least 30 minutes.

Galectin-8 staining was based on the method described by Boyle and Randow[44]. Fixed cells were washed twice with PBS and then incubated in permeabilization/block (PB) solution (0.1% Triton-X 100, 1% bovine serum albumin in PBS) for 30 minutes at room temperature, and then stained with goat anti-human galectin-8 antibody (R&D Systems, AF1305) diluted in PB solution overnight at 4°C. Cells were then washed three times with PBS and stained with AlexaFluor488-conjugated donkey anti-goat IgG (ThermoFisher, A-11055) diluted in PB solution for one hour at room temperature. Cells were then washed three times and kept in PBS for imaging.

A Nikon A1R laser scanning confocal microscope fitted with a 20x Plan Apo 0.75 NA objective was used to generate 2048 × 2048 pixel 8μm z-stacks consisting of 1μm optical sections. Image acquisition was carried out using a galvano scanner, 488nm and 561nm lasers, and a GaAsP multi-detector unit. Maximum intensity projections were generated in NIS Elements (Nikon) and used to calculate the percentage of THP-1 cells containing mycobacterial foci labeled with galectin-8 per imaging field.

### CCF-4 Assay & Flow Cytometry

CCF-4 assay was conducted as previously described[28], with minor modifications. Briefly, 1 × 10^6^ THP-1 cells were seeded on 12-well tissue culture plates and treated with 33 nM PMA for three days. Cells were then washed with media and then infected with single-cell suspensions of tdTomato-expressing Wildtype, *ΔESX-1*, or *ΔmmpL7 M. marinum* at a MOI of 1 for 4 h at 33°C in EM medium. 24 hours post infection cells were stained with Fixable Viability Dye eFluor660 (eBioscience). Cells were then harvested and stained for 1h at room temperature with 8 μM CCF4-AM (Invitrogen) in EM medium supplemented with 2.5 μM probenecid. Finally, cells were fixed overnight at 4°C in 4% (wt/vol) paraformaldehyde. Cells were analyzed in an LSRFortessa II cytometer, using FACSDiva software (BD Biosciences). At least 40,000 events per sample were collected. Data were analyzed using FlowJo (Treestar). Permeabilization percentage among infected monocytes was calculated using a region defined by an increased 450 nm signal (indicating CCF4 dye cleavage and, therefore phagosomal permeabilization) relative to uninfected cells in the same sample. Events in this region were designated “permeabilized”. Percent permeabilization was then calculated as a ratio of events in the permeabilized field over total live and infected cells.

### Hemolysis Assay

Hemolysis assays were conducted as previously described[28]. Briefly, defibrinated sheep red blood cells (RBCs) (Fisher Scientific) were washed twice and diluted to 1% (vol/vol) in PBS. Mycobacteria were grown in complete 7H9 media at 33°C to OD_600_ ~1.5, washed twice with PBS, and resuspended to ~30 OD units/mL. 100 μL *M. marinum* were mixed with 100 μL 1% RBC in a micro-centrifuge tube, centrifuged at 5000 × g for 5 minutes, then incubated at 33°C for 2 hours. Pellets were then resuspended, centrifuged at 5000 × g for 5 minutes and the A405 measured of 100 μL supernatant. PBS treatment was used as a negative control (background lysis) and 0.1% Triton X-100 (Sigma) as a positive control (complete lysis). Hemolysis was calculated as percentage of detergent lysis after subtracting background lysis.

## Acknowledgements

We thank Dr. Keith Boyle for advice on galectin-8 staining, and T. Kaewen Dang for proofreading of the manuscript.

